# Examination of Lipid Distributions in Hydrogel-Expanded Mouse Brain Tissue Using Imaging Mass Spectrometry

**DOI:** 10.1101/2024.09.26.615181

**Authors:** Jacob M. Samuel, Tingting Yan, Zhongling Liang, Boone M. Prentice

## Abstract

Imaging is an essential tool in biological research, and imaging mass spectrometry uniquely provides a label-free approach with high molecular specificity. However, imaging mass spectrometry is limited in spatial resolution, which in turn limits the biological structures and processes that can be studied at small dimensions. Custom lens setups and altered optical paths have been used to shrink the diameter of the incident laser beam probe in matrix-assisted laser desorption/ionization (MALDI) imaging mass spectrometry to achieve high spatial resolutions (< 5 μm). However, these research-grade instruments are complex and expensive, making high spatial resolution imaging experiments unrealistic for the broader community. An alternative method for improving spatial resolution is through physical magnification of the substrate, which has been well established in the subfield of expansion microscopy (ExM). ExM leverages superabsorbent hydrogels for isotropic expansion of tissues and retention of analytes containing fluorescent tags. While typical ExM involves covalently anchoring the analyte of interest to the hydrogel network, lipid retention without anchoring has been recently demonstrated for imaging mass spectrometry. Herein, we demonstrate expansion imaging mass spectrometry (ExIMS) of expanded, whole brain tissue and examine lipid distributions in both positive and negative ion mode across multiple brain structures. A linear expansion factor of 4.5-fold is achieved and used to obtain high spatial resolution images of mouse brain cerebellum. Approximately 95% of lipids in both positive and negative ion mode are retained in expanded tissue compared to unexpanded tissue. Additionally, the majority of lipid distributions across the brain are maintained post-expansion. Alterations to the hydrogel formulation (*e.g.*, crosslinker density) can significantly affect the ability of ExIMS to maintain accurate lipid distributions in expanded tissue.

## INTRODUCTION

Imaging mass spectrometry is a molecular imaging modality that is increasingly available on a variety of commercial mass spectrometry platforms.^1–3^ Despite the success and growing availability of imaging mass spectrometry, spatial resolution is limited to roughly 10-20 μm on most commercial platforms, and sub-cellular resolutions using advanced instrumentation have only rarely been achieved.^4, 5^ For example, in matrix-assisted laser desorption/ionization (MALDI) imaging mass spectrometry, spatial resolution is in part determined by the diameter of the incident MALDI laser beam.^6–9^ Newer instrument platforms have reported beam diameters down to 5 μm^10–12^ and recent research efforts have used custom lens setups,^13–16^ secondary lasers,^17, 18^ alternative laser wavelengths,^19–21^ altered optical paths,^22–27^ and modified sample stages^28^ to achieve beam diameters of 1-5 μm. However, research-grade instrument modifications are complex and expensive, making high spatial resolution imaging experiments unfeasible for the broader community. Even these setups begin to run into the diffraction limit, which is the theoretical limit to which the beam can be focused based on the laser wavelength. Operating at small beam diameters also introduces other challenges to the imaging mass spectrometry experiment, including the accuracy of the MALDI stage and the sensitivity of the measurements. For example, the step size of commercial MALDI stage motors is typically 5-10 μm, and operating at stage raster step sizes near or below this physical limit results in significant errors in step precision, distorting the image.^9, 29^ Many important biological processes occur within or between cells in spatially defined regions smaller than 10 μm (*i.e.*, roughly the average diameter of a mammalian cell), necessitating improvements in spatial resolution for imaging mass spectrometry workflows.

A novel, instrument-independent method to increase spatial resolution is via physical magnification of the tissue substrate. Previous attempts at “tissue stretching” have been performed by mounting thin tissue sections onto Parafilm membranes.^30, 31^ Manually pulling the membrane in opposite directions resulted in up to 5-fold improvements in effective spatial resolution.^32^ However, this method may be susceptible to nonlinear and/or asymmetrical stretching in the x and y dimensions, poor reproducibility, and low sensitivity. Alternatively, controlled isotropic tissue expansion can be performed using a hydrogel polymer network. This framework has been established in expansion microscopy (ExM), in which fluorescent dyes conjugated to antibody labels are anchored to a polyelectrolyte hydrogel.^33^ Upon exposure of the anchored system to an aqueous environment, the hydrogel network swells in size, physically magnifying the tissue up to ∼20-fold. The original ExM method infused a sodium acrylate/acrylamide hydrogel with N,N-methylenebisacrylamide as a cross linker throughout a 30 µm mouse brain section.^34^ The tissue is then digested with proteinase K to homogenize mechanical characteristics and ensure isotropic expansion upon submersion in doubly deionized (DI) water, which results in ∼4.5-fold linear expansion. The target protein is retained through digestion and expansion due to the use of a trifunctional oligonucleotide that has a methacryloyl group that can take part in free radical polymerization, linking the protein to the hydrogel and a fluorophore for visualization. The evolution of ExM over the past nine years has altered these steps for a wide variety of applications. Some notable developments are expansion for different analytes,^35–37^ expansion of formalin-fixed paraffin embedded samples,^38^ adjusting the gel composition for increased fidelity^39^ or expansion factor,^40–42^ and adjusting the digestion to expand a variety of sample types.^43–45^

Expansion-based sample preparation was only recently shown to be viable for imaging mass spectrometry analysis, with novel reports showing success in mouse brain, liver, and testis tissue.^46–48^ These publications have all featured lipid imaging, which has only been performed in conventional ExM via labeling or anchoring of the lipids to the hydrogel.^49–51^ Lipid imaging mass spectrometry expansion methods do not require covalent linking to the gel, indicating lipids can instead be physically retained by the polymer network. Due to the complex sample preparation, ensuring minimal delocalization and maintaining accurate lipid distributions are crucial to performing reliable expansion imaging mass spectrometry (ExIMS).^52, 53^ Herein, we describe an ExIMS protocol and investigate the mechanism of lipid retention by characterizing the lipids retained in the gel, comparing their distributions between expanded and unexpanded serial tissue sections, and assessing how adjustment of the expansion protocol affects these distributions. Our work also highlights optimized polymer designs and computational methods that lead to improvements in lipid retention and monitoring spatial accuracy.

## EXPERIMENTAL

### Chemicals and Reagents

HPLC-grade water and sodium chloride were obtained from Fisher Chemical (Waltham, MA, USA). 1,5-diaminonaphthalene (DAN), 4-hydroxy-2,2,6,6-tetramethylpiperidin-1-oxyl (4HT) ammonium persulfate (APS), N,N’-methylenebisacrylamide(MBAA), acrylamide and tetramethylenediamine (TEMED) were purchased from Sigma-Aldrich (St. Louis, MO, USA). Proteinase K was purchased from New England Biosciences (Ipswich, MA, USA). Mouse brain (C57BL/6) tissue was acquired from BioIVT (Westbury, NY, USA). 4% paraformaldehyde was acquired from Boster Bio (Pleasnton, CA, USA). Sodium acrylate and 9-aminoacridine and were purchased from Santa Cruz Biotechnology (Dallas, TX, USA). Acryloyl-X (6-((acryloyl)amino)hexanoic acid, succinimidyl ester)was purchased from ThermoFisher Scientific (Waltham, MA, USA).

### Tissue Preparation

Tissue sectioning was performed on a Leica CM3050 S Cryostat (Leica Biosystems, Wetzlar, Germany). Mouse brain was sectioned to 30 μm thickness and then thaw mounted onto either a microscope slide for ExIMS or an indium-tin oxide (ITO) slide for normal imaging mass spectrometry analysis. Sections were taken in pairs with the first being expanded and the serial section acting as an unexpanded comparator. Tissues were stored at -80 °C.

### ExIMS

#### Gelation

Tissue sections were covered with a 4% paraformaldehyde solution and incubated at 4 °C for 15 minutes. The tissues were then washed three times with 1x phosphate buffered saline (PBS). Tissues were then coated in an acryloyl-X solution (0.3 mg/mL in 1x PBS) at room temperature for at least 6 hours, covered. The tissues were washed twice with 1x PBS, for 10 minutes each. Tissue sections were incubated in the monomer solution (1x PBS, 2 M NaCl, 8.625% (w/w) sodium acrylate, 2.5 % (w/w) acrylamide, 0.15 % (w/w) MBAA, 0.01 % (w/v), 4HT, 0.2% (w/v) APS and 0.2% (w/v) TEMED for 30 minutes at 4°C. Prior to embedding, the monomer solution was cooled to 4°C to prevent premature gelation. Tissue sections, still in monomer solution, were enclosed in a constructed gelling chamber, as described previously,^34^ and allowed to sit at room temp for 3.5 hours.

#### Digestion

Tissues embedded in hydrogels were cut out of incubation chambers and excess gel was removed. The sample was submerged in proteinase K solution (4 U/L in 50 mM Tris, 0.8 M NaCl, pH 8 buffer) at room temperature overnight.

#### Expansion and Sample Mounting

The digestion buffer was removed and the sample was washed three times with cold 1x PBS and then submerged in excess de-ionized H_2_O. The sample remained in the ddH_2_O until the desired or max expansion was achieved, with the water being exchanged every twenty minutes. Expanded tissues were cut down to the appropriate size, flipped, and carefully transferred to an ITO slide using a coverslip. Tissues were then desiccated in a vacuum desiccator under pressure at 2 Torr until the height of the gel appeared to reach a minimum height evenly across whole sample. The tissues then remained in a desiccator overnight to further remove any moisture.

### Imaging Mass Spectrometry

A DAN MALDI matrix layer was applied to desiccated samples using an M5 TM sprayer (HTX Technologies, LLC, Chapel Hill, NC, USA). The spraying solution consisted of a 10 mg/mL of DAN matrix solution in 90% acetonitrile and 10% water. The sprayer temperature was set to 30°C, with a flow rate of 0.1 mL/min, spray velocity of 1200 mm/min, track spacing of 2.5 mm, and a pressure of 10 psi. Five passes of the matrix were applied resulting in ∼2 mg of matrix applied to the tissue and microscope slide. Imaging mass spectrometry experiments were performed on a SolariX XR 7T FT-ICR mass spectrometer (Bruker Daltonics, Billerica, MA) equipped with an Apollo II dual MALDI/ESI source and Smartbeam II Nd:YAG laser system (355 nm, 2kHz frequency). Positive ion mode lipid imaging was conducted using 30-50 laser shots at 27% power on a 50% global attenuation setting. Negative ion mode lipid imaging was conducted using 100-150 laser shots at 32% power on a 50% global attenuation setting (note that the number of laser shots was adjusted to generate the same approximate base peak intensity between expanded and unexpanded tissues). High resolution accurate mass (HRAM) spectra were acquired using a resolving power of ∼270,000 at *m/*z 760 via a ∼4 second transient. External mass calibration was performed using red phosphorus and tentative lipid identifications were made using a <5 ppm mass accuracy threshold.

### Microscopy

All microscopy was performed using an Axio Imager M2 brightfield/fluorescence microscope and slide scanner (Carl Zeiss, Jena, Germany). H&E staining was performed on unexpanded mouse brain serial sections following imaging mass spectrometry analysis by first washing off the matrix layer and then staining, as described in previous literature.^54^ Autofluorescence images were acquired pre and post-expansion with excitation at 385 nm and emission 465 nm.

### Data Analysis

Images were processed using fleximaging. Lipid identification was performed by searching previous reports and the LIPIDMAPS database.^55–57^ Image registrations and comparisons were performed in MATLAB. Residual values were calculated with the following equation:

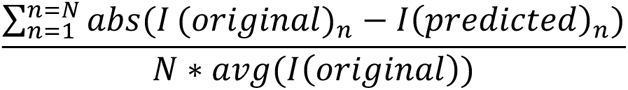

where N is the total number of pixels, I(original)_n_is the intensity value of the n^th^ pixel in the unexpanded tissue mass spectrometry image, and I(predicted)_n_ is the intensity value of the n^th^ pixel in the expanded mass spectrometry image. Expansion factors were calculated by comparing distances between pairs of easily seen features in autofluorescence images of the sample pre- and post-expansion in multiple regions and orientations of the tissue, and taking the average across all of the measurement pairs.

## RESULTS

### ExIMS Workflow

The ExIMS workflow described herein is based off of original ExM workflows and recent reports on applying tissue expansion to imaging mass spectrometry (**Figure 1**).^46–48^ Briefly, the workflow begins with fixation of the mouse brain sections followed by incubation with AcX, which converts primary amines in molecules throughout the tissue into acrylamide (**Figure 1a**). This allows the tissue to be anchored to the hydrogel during free radical polymerization (**Figure 1b**), and thus expand isotropically with the swelling of the hydrogel. Free radicals are generated by APS, modulated by 4HT and TEMED, creating polymer chains with monomers of acrylate and acrylamide. MBAA acts as a crosslinker, linking polymer chains together and providing 3-D structure and support for the gel. The tissue is then digested (**Figure 1c**), in this case using proteinase K, to break down structural proteins, allowing for isotropic expansion of the tissue without tearing. Following digestion, the tissue can be expanded via submersion in ddH2O (**Figure 1d**), in which water is drawn into and retained in the hydrogel by a combination of osmosis due to ionic species, the hydrophilicity of polymer components, and hydrogen bonding with the polymer side chains. Expansion is also induced by removal of sodium ions associated with the acrylate groups during washing steps, resulting in a negatively charged polymer network that electrostatically repels chains away from each other.^58–60^ The expanded tissue is then flipped (*i.e.*, so the tissue side is facing up for analysis) and transferred to an ITO slide using a coverslip before undergoing desiccation to flatten the gel (**Figure 1e**). Passive desiccation in a typical drying cabinet produced inconsistent results, often resulting in tears or excessive shrinking. These issues were also noted during vacuum desiccation when an excess of moisture was present (*e.g.*, when attempting to dry a large number of samples simultaneously). Desiccation via a vacuum desiccator at lower pressures (*i.e.*, 1-2 torr), produced consistent desiccation of the hydrogels without tearing or shrinking. Post-desiccation, conventional imaging mass spectrometry sample preparation (**Figure 1f)** and analysis (**Figure 1g**) is performed. Modifications in our ExIMS workflow compared to previously reported methods include a longer fixation time and a room temperature polymerization and digestion steps. Fixation with paraformaldehyde is generally light to prevent tearing and loss of tissue during the extensive sample preparation. However, our experiences with shorter fixation times result in tissue that is visibly lost during the digestion step due to the fragility of the sample. Additionally, polymerization was conducted at room temperature (compared to conventional 37 °C) and for a longer time (*i.e.*, 4 hours here compared to 2 hours) in order to reduce the fluidity of lipids at higher temperatures so as to better maintain accurate lipid distributions.

**Figure 1.**
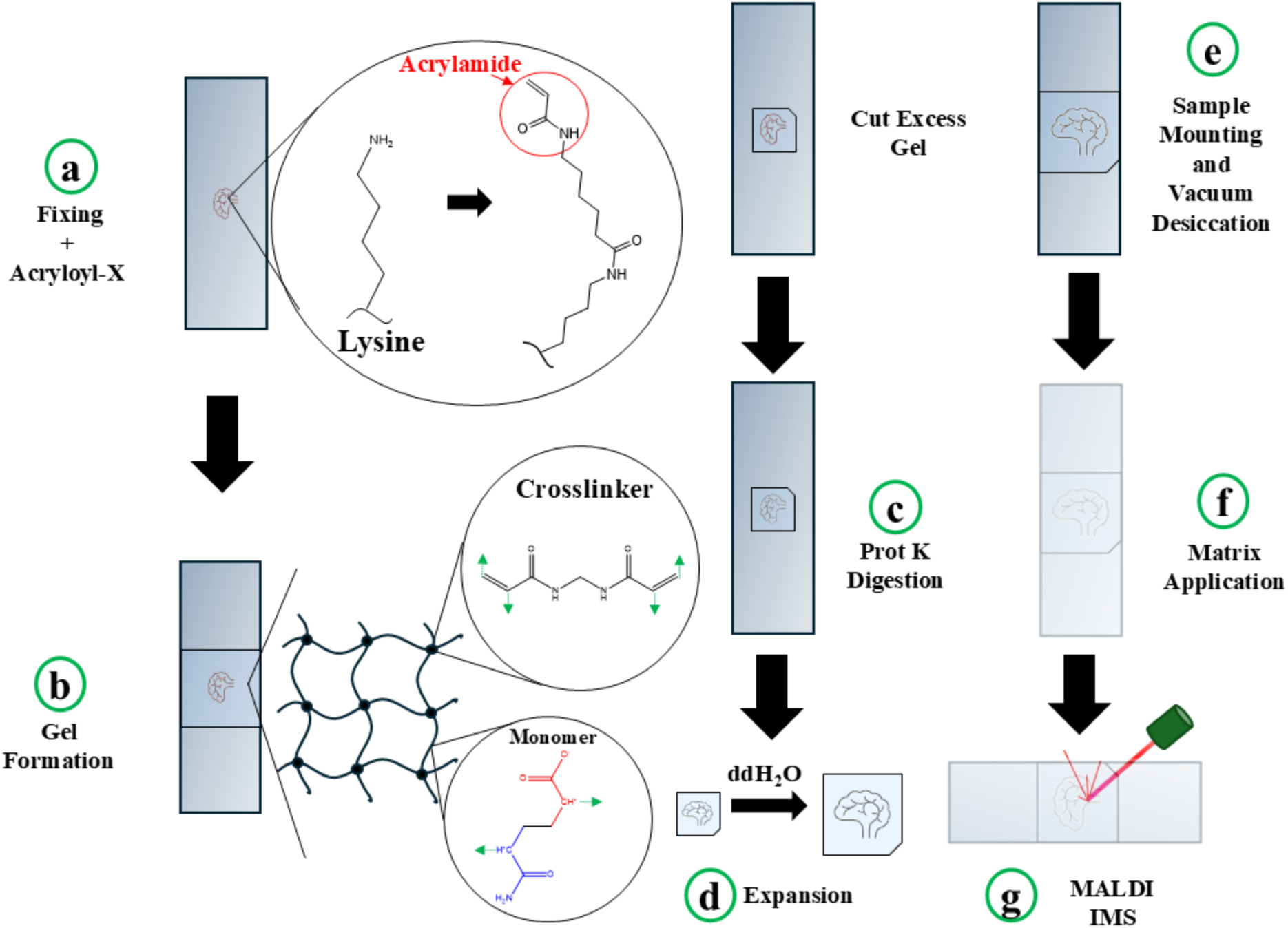
An overview of the ExIMS workflow: (a) Samples are first fixed and then incubated in an acryloyl-x solution that converts primary amines into polymerizable acrylamides. (b) Samples are then infused with a monomer solution consisting of acrylamide (blue) and acrylate (red) as monomers and N’N’-methylene(bis)acrylamide for crosslinking, and then incubated at room temperature to undergo free radical polymerization. (c) The sample is then digested to mechanically homogenize the sample to ensure even expansion. (d) The sample is then submerged in ddH_2_O to enable expansion. (e) The sample is desiccated under vacuum and (f) matrix is applied using a robotic sprayer. (g) Finally, the sample is analyzed via imaging mass spectrometry.

This methodology was used to improve the effective spatial resolution in both positive and negative ion mode lipid imaging mass spectrometry analysis of mouse brain tissue. A portion of mouse brain cerebellum was expanded approximately 4.5-fold and analyzed using a 30 µm raster width in positive ion mode (**Figure 2).** Juxtaposed with imaging mass spectrometry analysis of the same region from an unexpanded serial tissue section acquired at the same raster width (**Figure 2c**), the increase in effective spatial resolution is evident in the expanded tissue (**Figure 2b**). Purkinje cells and various cell layers of the cerebellum are clearly visible in the expanded gel and confirmed via registration with a histological image (**Figure 2a**). The tissue expansion produces an image that is effectively equivalent to an image acquired at roughly 6.6 µm pixel size (*i.e.*, 30 µm raster step divided by a 4.5-fold linear expansion), which corresponds to a roughly 20-25-fold increase in pixel density. While there was some decrease in lipid signal intensity in the expanded tissue due to dilution, the effect was minimal and not proportional to the expanded volume. This required only minor adjustments to the number of laser shots used to acquire the images (*i.e.*, 40 laser shots for unexpanded tissue and 50 laser shots for expanded tissue). Signal loss is likely mitigated by the high abundance and high ionization efficiency of the detected phosphatidylcholines (PC). Additionally, potassiated PC ion types are not observed in the expanded tissue. Potassium ions are washed away during the expansion sample preparation, though sodium ions are retained as a part of the initial hydrogel formulation. The lack of potassiated ion types helps condense PC ion signal into only the protonated and sodiated ion type channels.

**Figure 2.**
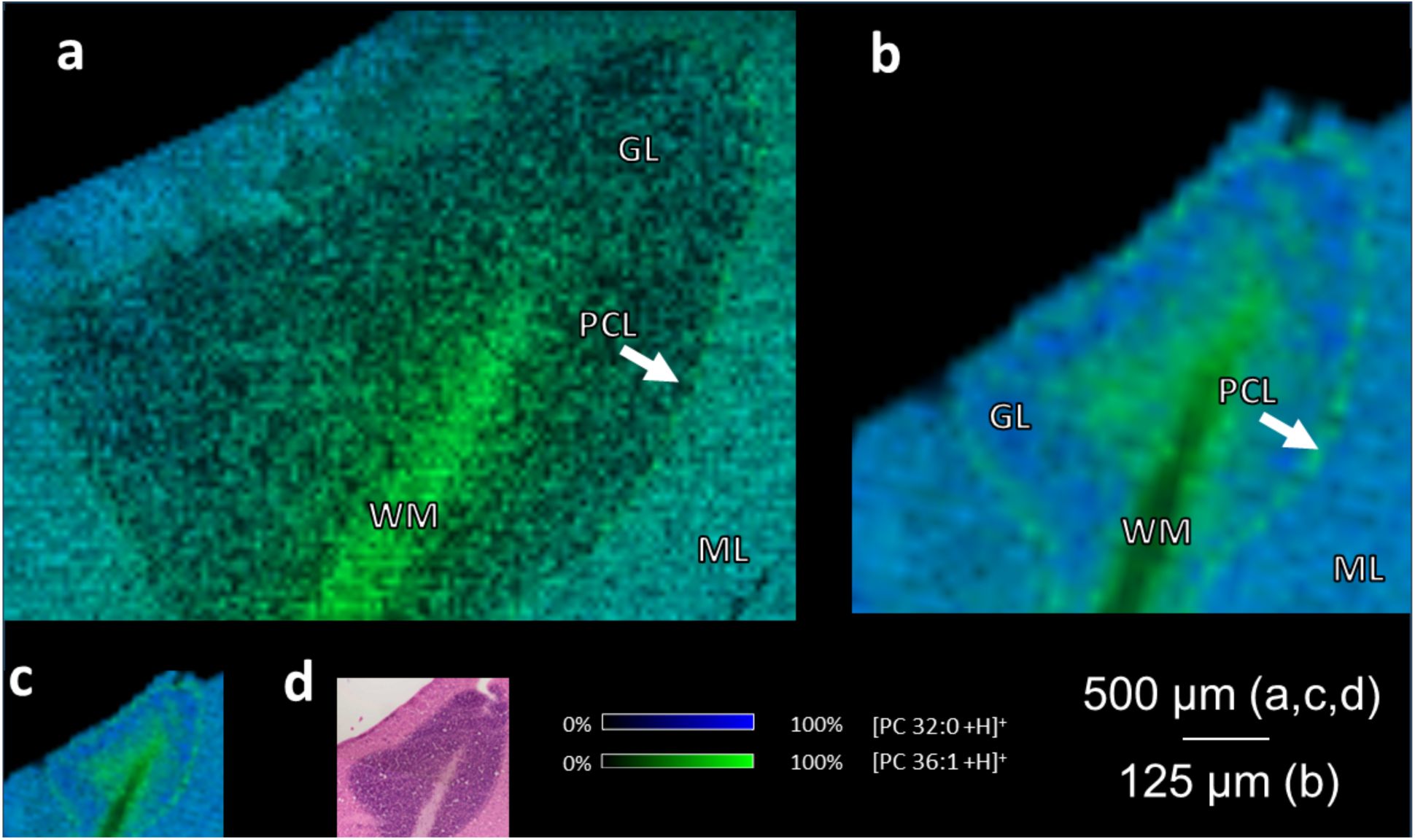
(a) Mouse brain cerebellum is analyzed via positive ion mode imaging mass spectrometry on (a) expanded and (b, c) unexpanded tissue sections. Imaging mass spectrometry is performed using a 30 μm raster step in both analyses, resulting in approximate effective spatial resolutions of 6.6 μm and 30 μm in the expanded and unexpanded tissue sections, respectively. PC 36:1 and PC 32:0 are localized to regions of the cerebellum (ML = Molecular Layer, GL = Granular Layer, PCL = Purkinje Cell Layer, WM = White Matter). (d) H&E staining was performed on the unexpanded serial section following imaging mass spectrometry.

High mass resolution spectra (**Supplemental Figure S1**) acquired on various regions of expanded and unexpanded tissues enable the identification of 85 lipids in positive ion mode and 158 lipids in negative ion mode (**Supplemental Table S1**). The number of lipids identified in expanded versus unexpanded tissues show remarkable similarities in both negative and positive ion mode, with only ten additional lipids identified in the unexpanded tissues for both ionization polarities (**Figure 3**). Similar classes of phospholipids are detected in unexpanded and expanded tissue, indicating broad lipid retention during expansion. Surprisingly, lipids with primary amines, which are potential targets of both the fixing and anchoring steps during cross-linking, are detected in comparable quantities between the tissue types. For example, six more PE compounds were identified in the unexpanded tissue in positive ion mode and three more in negative ion mode. Formaldehyde has been shown as an ineffective crosslinker for lipids, and lipid crosslinking experiments typically require more specialized reagents.^61–63^ While there is some loss, PE and PS retention may also be due to the high abundances of these lipids, and the negatively charged inner leaflet of the plasma membrane, which contains the majority of PE’s and PS’s, being non-conducive to these reactions..^64^ The ability to retain a wide variety of lipids in both ionization polarities is promising for future expansion experiments and suggests a universal retention mechanism not related to specific chemical interactions with lipid headgroups, but rather more general characteristics such as size and lipophilicity. Notably, this comparison of lipid retention is for untreated and unexpanded tissue compared to expanded tissue, which undergoes several washes in PBS, proteinase K buffer (pH 8 tris buffer), and water during the sample preparation. Prior literature indicates that brief tissue washing with an aqueous solvent can remove endogenous salts and improve lipid limits of detection in positive ion mode,^65^ which may mitigate some of lipid signal dilution due to expansion, as noted above.

**Figure 3.**
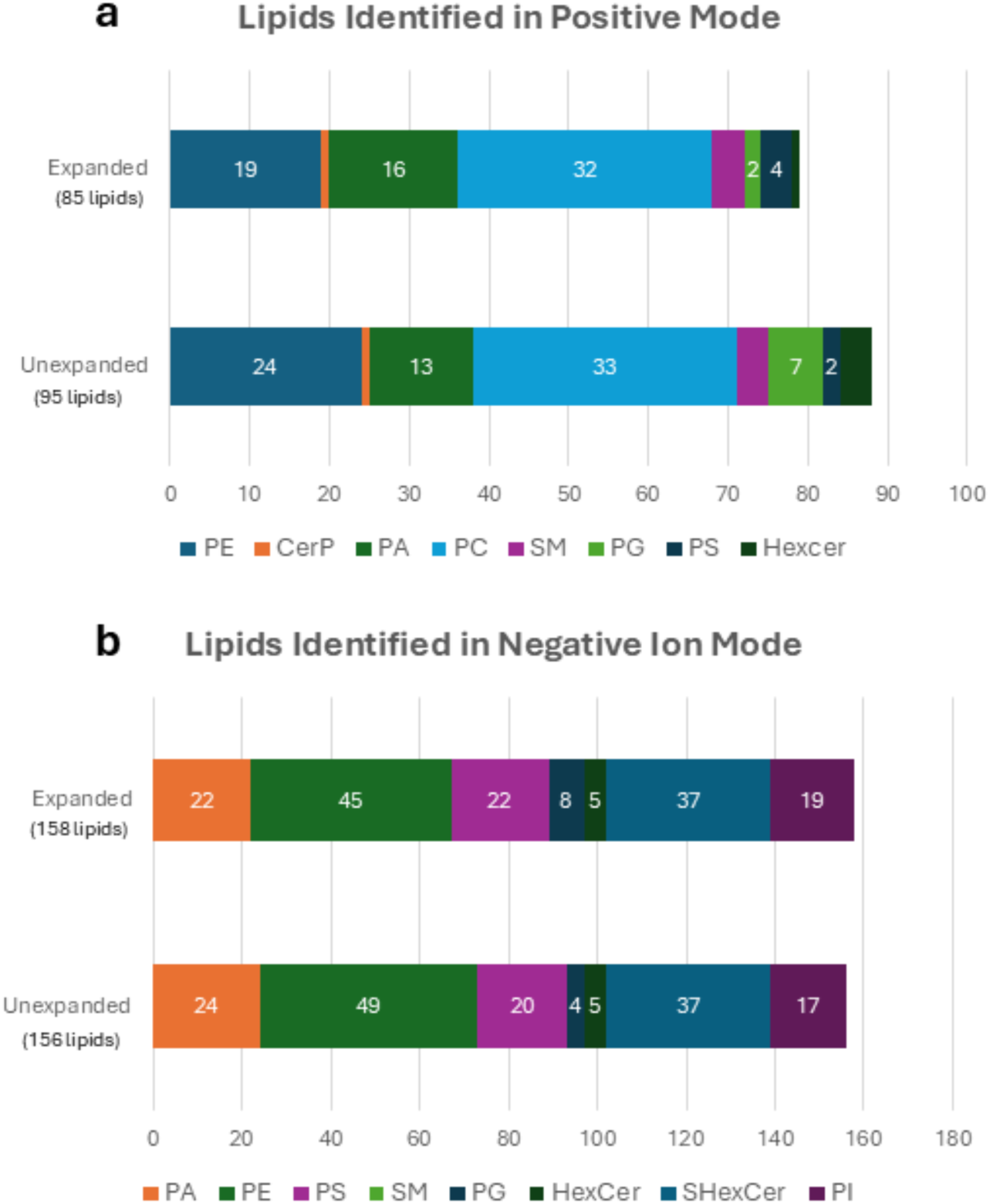
Number of lipids identified in (a) positive and (b) negative ion modes in both expanded and unexpanded mouse brain tissue.

### Characterizing Expansion Accuracy

The extent of tissue distortion during the expansion process was characterized via microscopy and registration comparisons between unexpanded and expanded serial tissue sections (**Figure 4**). Imaging mass spectrometry of expanded and unexpanded serial tissue sections was first acquired at the same raster step size. Ion images in the two experiments were co-registered using non-rigid registration in MATLAB to facilitate spatial comparisons. H&E staining was also performed on unexpanded tissue sections following imaging mass spectrometry to provide a high spatial resolution image to confirm tissue morphology. Pearson correlation coefficients (PCC) were calculated by comparing the pixel-by-pixel intensity values for the co-registered unexpanded and expanded images. PCC scores closer to 1 indicate a stronger correlation of spatial patterns between the two images. An evaluation of protonated sphingomyelin (SM) d36:1 (*m/*z 731.6072, 1.47 ppm) revealed a PCC score of 0.78 (**Figure 4a**). The PCC score indicates that the lipid distribution was maintained fairly accurately between the two tissue sections, especially considering that the tissue sections are serial and thus expected to differ somewhat in their spatial similarity. Additionally, the difference in effective spatial resolutions between the two images will also contribute to dissimilarity. The overlaid image highlights regions with differences in signal intensity, with green indicating regions where the expanded tissue had higher relative intensity and purple highlighting regions of higher relative intensity in the unexpanded image (**Figure 4a**). Similar spatial analyses were conducted on smaller regions of the brain using different lipid ions. For example, a comparison of PC 34:1 (*m/z* 760.5850) in the cerebellum between the co-registered expanded and unexpanded serial tissues results in a PCC score of 0.89 (**Figure 4b**).

**Figure 4.**
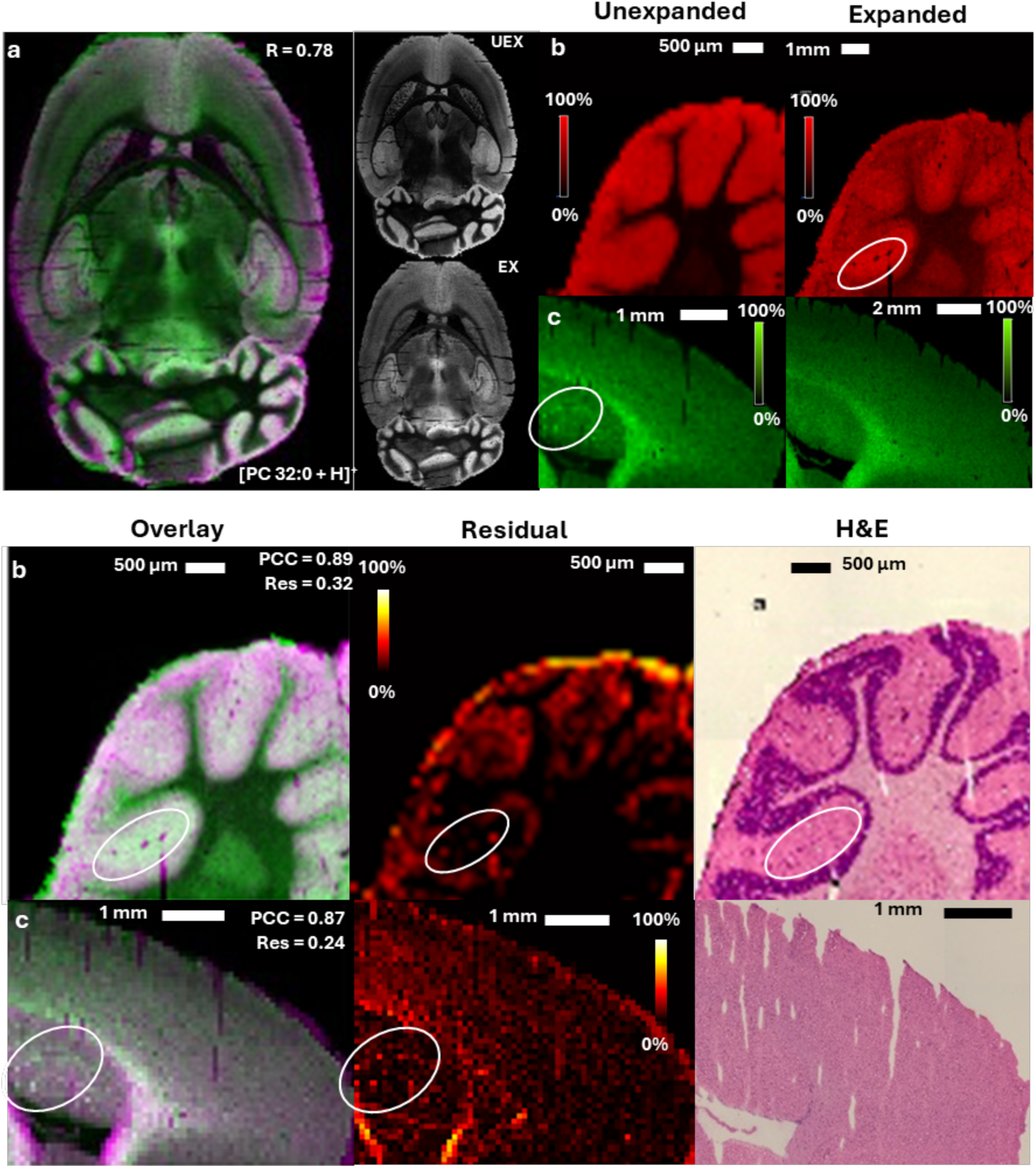
Co-registration and comparison of expanded and unexpanded tissue. (**a)** An overlay of the ion images for [PC 32:0 + H]^+^ from expanded and serial unexpanded mouse brain sections shows areas of greater signal intensity in the expanded tissue (green) and areas of greater signal intensity in the unexpanded tissue (purple). Co-registered images of the unexpanded (top) and expanded (bottom) tissues are shown in gray scale. Lipid spatial distributions for (**b)** [PC 34:1 + H]^+^ in the cerebellum and (**c)** [PC 36:1 + H]^+^ in a portion of the corpus callous and cortex are compared for (from left to right and continuing below) unexpanded tissue, expanded tissue, an overlay of the co-registered images, the residuals image, and an H&E stain of the unexpanded serial section following imaging mass spectrometry.

Relative average residual values were also calculated using the average of the absolute value of the intensity differences between the expanded and unexpanded images. Smaller relative residual values indicate greater similarity in ion intensities. The relative average residual value for PC 34:1 in the cerebellum is 0.32, reflecting high similarity in the intensity patterns between the pre- and post-expanded ion images. Residual images were generated to analyze spatial similarity across the tissue sections by plotting the absolute differences between the intensity values of the unexpanded image and those in the expanded image. Higher intensities in a residual image indicate areas of greater discrepancy, which is observed with PC 34:1 in the granular layer and in a tissue fissure (circled in white in **Figure 4**). Still, the layers of the cerebellum are resolved with greater clarity in the expanded tissue, and are confirmed in the H&E imaging. The same analysis was also performed for PC 36:1 (*m/z* 788.6166, 0.29 ppm) in the corpus callosum (**Figure 4c**), yielding a PCC value of 0.87. Here, the residual image shows high similarity between the images, with minor dissimilarity in the striatum. However, a pattern of striosomes (circled in **Figure 4c**) is visible in the unexpanded tissue but not the expanded brain.

Spatial distributions for lipids identified in both positive and negative ion modes were examined across entire mouse brain tissue sections. Samples were expanded with a 2-fold linear expansion factor to allow for imaging the entire expanded brain in one experiment (*i.e.*, a 4.5-fold expansion results in a mouse brain section too large to fit on a single microscope slide and also requires a substantially longer imaging analysis time).^10^ Spatial distributions for most lipids across the whole mouse brain are generally maintained (**Figure 5**), with minor exceptions. As a result of repeated washing steps and the presence of sodium ions in the hydrogel formulation, the biological relevance from sodium-adducted ion types is lost. The spatial distributions of sodiated lipid ion types in expanded tissue instead mimic distributions of the protonated ion types in the unexpanded tissue (**Supplemental Figure S2**). One region of spatial dissimilarity is in the cerebellum, where some lipids do not maintain sharp boundaries between the white matter and granular layer (*e.g.*, [PI 38:4 - H]^−^ and [PC 38:6 + H]^+^). Other minor dissimilarities are detectable in the thalamus and striatum, but strong similarity is observed for most lipids.

**Figure 5.**
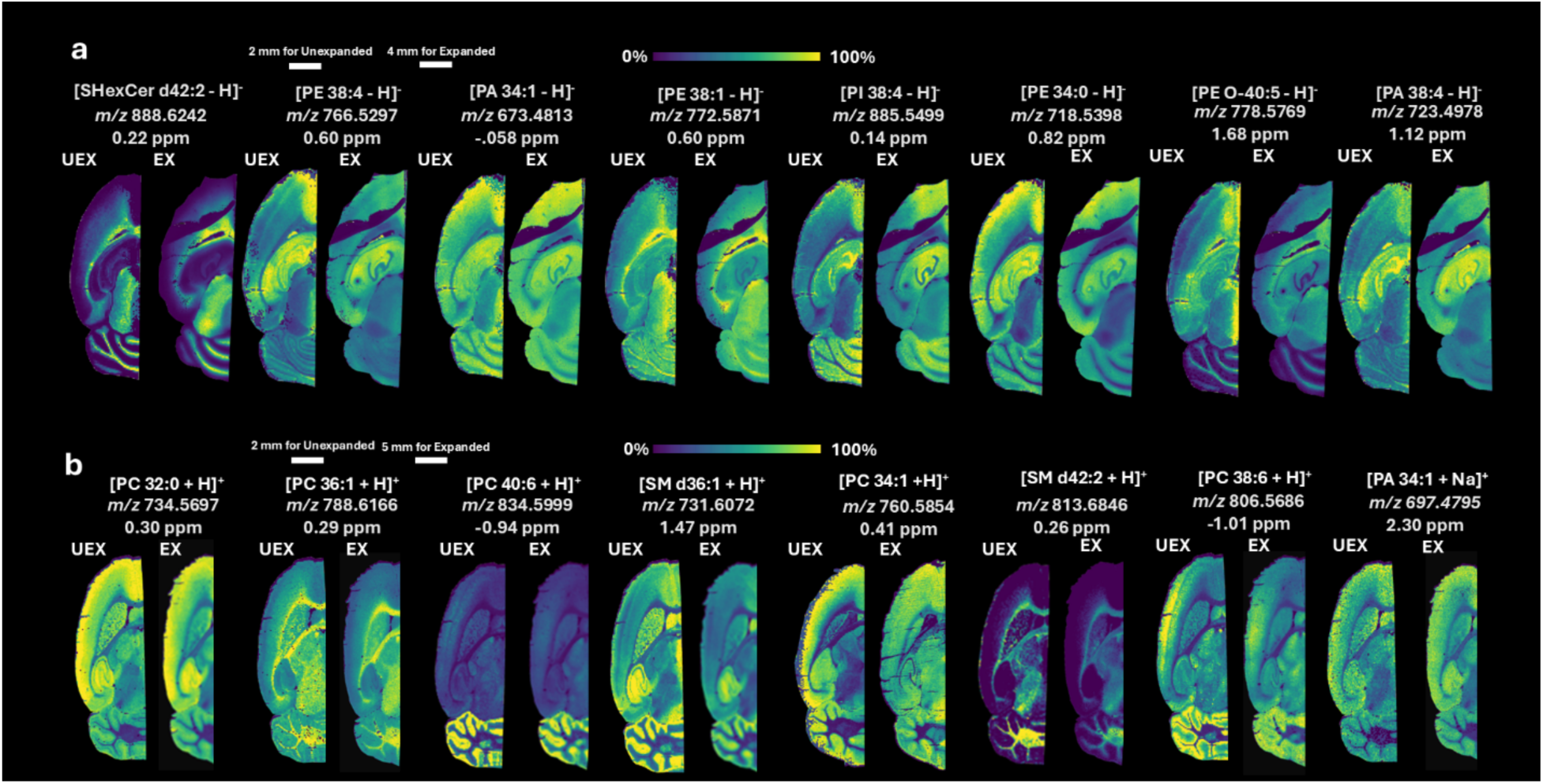
Spatial distributions of lipids detected in (a) positive and (b) negative ion modes are largely similar in serial expanded and unexpanded mouse brain tissue sections. A 2-fold linear expansion factor was used for expanded tissues and all images were acquired using an 80 μm raster step. Note that expanded tissues are 2-fold larger (as is evident by the respective scale bars), but have been formatted to the same size as the unexpanded tissue to enable comparisons in lipid distributions.

Dissimilarities in lipid retention in these regions may arise from differences in the tissue environments of these substructures (*e.g.*, densities, lipid concentration). However, it is possible to alter the hydrogel formulation and polymerization steps to better maintain lipid distributions. For example, the temperature during polymerization and the crosslinker density were adjusted during sample preparation to alter lipid retention during ExIMS (**Figure 6**). Gels were polymerized either at 37 °C (as is common is most expansion protocols) or at room temperature. Crosslinker density was adjusted by changing the amount of MBAA in the monomer solution, with more MBAA resulting in a higher crosslinker density The resulting images of the cerebellum were compared visually and by using PCC and relative average residual calculations. The PCC and residual values calculated for ion images of PC 36:1 show little difference across the different sample preparation conditions, which is consistent with this lipid maintaining similar distributions pre- and post-expansion (*i.e.*, high abundance in the white matter) (**Figure 6a**). By comparison, PC 32:1 generally shows dissimilar spatial distributions post-expansion compared to unexpanded tissue (**Figure 6b**). Only the high crosslinker preparation produces a spatial distribution with high similarity to the unexpanded tissue (*i.e.*, high abundance in the granular layer and lower abundance in the white matter). The three other expansion formulations instead show similar abundances in both the white matter and granular layer, with no visible boundary. This difference is reflected in the calculated relative average residual values for the PC 32:1 ion images, which are generally 5-10-fold larger than the residual values calculated for the PC 36:1 ion images, except for the high crosslinker formulation, which is only roughly 2-fold greater. Notably, the difference in residual value is greatest for the low crosslinker formulation. This is consistent with qualitative evaluation of the other expansion formulations, which contain lower signal intensity in the white matter and higher signal intensity in the granular layer, while the low crosslinker formulation shows the opposite distributions (*i.e.*, higher signal intensity in the white matter and a clear band of lower signal intensity surrounding this in the granular layer as indicated by the arrows) (**Figure 6b**). This further suggests that crosslinker density can play a critical role in retaining accurate lipid distributions. This general trend for gel formulation is consistent for other lipids that have similar distributions in the granular layer, such as PC 36:4 and PC 38:6 (**Supplemental Figure S3**).

**Figure 6.**
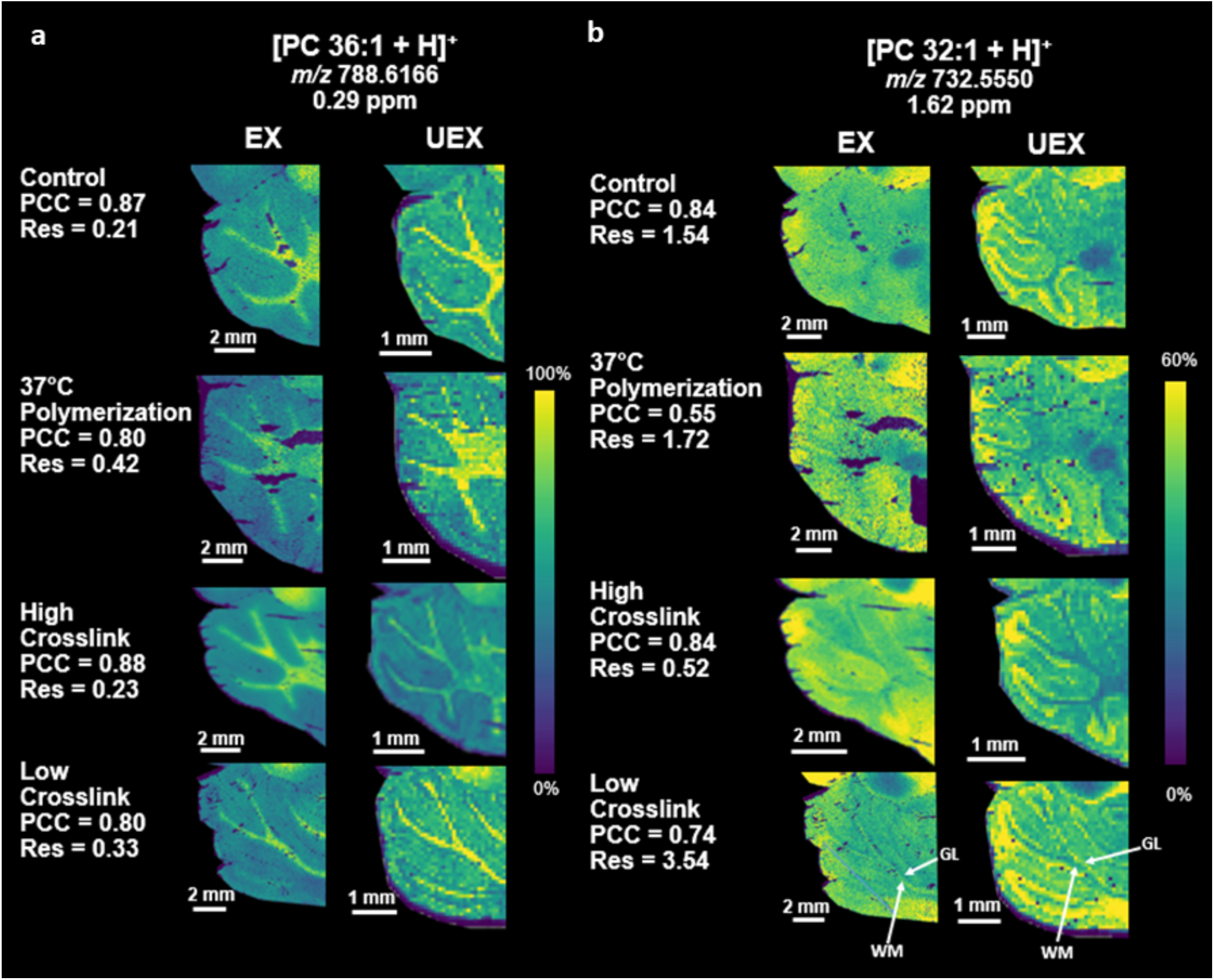
ExIMS of mouse brain sections is performed using variations in the expansion procedure, including polymerizing at 37 C° for 2 hours versus at room temperature for 4 hours and either doubling or halving the crosslinker concentration in the monomer solution. PCC and relative average residual scores are used to compare levels of similarity between expanded (left) and unexpanded (right) imaging mass spectrometry analyses of (a) [PC 36:1 + H]^+^ and (b) [PC 32:1 + H]^+^. The PCC and residual values correspond to comparisons between the expanded and unexpanded serial tissue sections after non-rigid registration. Arrows indicate the granular layer (GL) and white matter (WM).

The effect of polymerization temperature on lipid retention was also briefly examined, with the thought that sub-ambient temperatures may reduce lipid fluidity and therefore better preserve lipid distributions. However, no differences were observed between the two methods (**Supplemental Figure S4**). Lipid detection was not significantly affected by adjustment of temperature during the expansion protocol, though thermally labile analytes may benefit from a more ambient expansion procedure. This suggests that altering gel formulations (*e.g.*, increasing crosslinker density) can provide for more accurate ExIMS results, though it should be noted that some minor dissimilarities persist in the current formulation (*e.g.*, in the previously mentioned striatum), requiring further refinement of the expansion procedure. Still, increasing the crosslinker density likely better retains lipids through physical polymer entanglement. However, this may limit on ultimate expansion factor that can be reached for polymer-entangled analytes in a single gel, given that expansion requires the polymer strands to stretch apart as volume increases. This is in contrast to covalently anchored analytes or iteratively expanded samples used in other ExM approaches.

## CONCLUSIONS

We have described an expansion imaging mass spectrometry (ExIMS) workflow for lipid analysis in mouse brain that provides for a 4.5-fold increase in effective spatial resolution. This method provides a instrument-independent manner to increase imaging mass spectrometry spatial resolution without the need for specialized laboratory instrumentation. While shown here for MALDI imaging mass spectrometry, this sample preparation method is applicable to other imaging modalities (*e.g.*, desorption electrospray ionization [DESI] and secondary ion mass spectrometry [SIMS]) and is readily applied to any commercial imaging MS instrument platform. ExIMS requires an intensive sample preparation procedure that requires care to maintain the integrity of the fragile tissue embedded in the hydrogel. ExIMS protocols also likely need to be adjusted when analyzing new tissue types and analytes. The lipid content following hydrogel-based tissue expansion was identified here using HRAM measurements in both positive and negative ion modes, showing that roughly 95% of lipids are retained following expansion. The accuracies of lipid spatial distributions following expansion were determined using non-rigid registration and calculation of PCC and residual scores to determine spatial similarity. Overall, ExIMS shows minimal tissue distortion and good spatial correlation, allowing for confirmation of small structures in expanded tissue with high confidence. Variations in the tissue expansion protocol can affect the resulting lipid distributions. For example, higher crosslinker density can better retain lipids during expansion and results in more accurate spatial distributions in small brain substructures. These results suggest that further refinement of ExIMS protocols may allow for more rigorous and expanded tissue analyses.

## ACKNOWLEDGEMENTS

This work was supported by a generous contribution from Eli Lilly and Company.

## Notes

### Competing Interest Statement

The authors have declared no competing interest.

