## Supplemental Information for "Examination of Lipid Distributions in Hydrogel-Expanded Mouse Brain Tissue Using Imaging Mass Spectrometry"

Jacob M. Samuel, Tingting Yan, Zhongling Liang, Boone M. Prentice*
Department of Chemistry, University of Florida, Gainesville, FL 32611

*Address correspondence to:
Dr. Boone M. Prentice
214 Leigh Hall
PO Box 117200
Department of Chemistry
University of Florida
Gainesville, FL 32611, USA


Key words: imaging mass spectrometry, MALDI, high resolution, expansion, hydrogel

**Supplemental Table 1:** Lipids tentatively identified using HRAM measurements in positive and negative ion mode analysis of fresh frozen mouse brain tissue.

| Observed m/z | Theoretical m/z | mass error | species | ion | formula |
|---|---|---|---|---|---|
| 622.4450 | 622.4442 | 1.22 | PC 24:0 | [M+H]+ | C32H64NO8P |
| 650.4412 | 650.4391 | 3.19 | PE 28:1 OH | [M+H]+ | C33H64NO9P |
| 650.4785 | 650.4755 | 4.61 | PC 26:0 | [M+H]+ | C34H68NO8P |
| 672.4233 | 672.4211 | 3.27 | PE 28:1 OH | [M+Na]+ | C33H64NO9PNa |
| 673.5162 | 673.5167 | -0.73 | PA O-35:2 | [M+H]+ | C38H73O7P |
| 676.4556 | 676.4548 | 1.22 | PE 30:2 OH | [M+H]+ | C35H66NO9P |
| 678.4725 | 678.4704 | 2.97 | PE 30:1 OH | [M+H]+ | C35H68NO9P |
| 694.5165 | 694.5146 | 2.68 | CerP d38:2 | [M+Na]+ | C38H74NO6PNa |
| 695.4642 | 695.4622 | 2.81 | PA 34:2 | [M+Na]+ | C37H69O8PNa |
| 697.4795 | 697.4779 | 2.30 | PA 34:1 | [M+Na]+ | C37H71O8PNa |
| 700.4541 | 700.4524 | 2.44 | PE 30:1 OH | [M+Na]+ | C35H68NO9PNa |
| 706.5400 | 706.5381 | 2.60 | PC 30:0 | [M+H]+ | C38H76NO8P |
| 719.4615 | 719.4622 | -1.04 | PA 36:4 | [M+Na]+ | C39H69O8PNa |
| 720.5551 | 720.5538 | 1.84 | PE 34:0 | [M+H]+ | C39H78NO8P |
| 721.4766 | 721.4779 | -1.84 | PA 36:3 | [M+Na]+ | C39H71O8PNa |
| 723.4946 | 723.4935 | 1.51 | PA 36:2 | [M+Na]+ | C39H73O8PNa |
| 725.5104 | 725.5092 | 1.67 | PA 36:1 | [M+Na]+ | C39H75O8PNa |
| 728.5217 | 728.5201 | 2.17 | PC 30:0 | [M+Na]+ | C38H76NO8PNa |
| 729.5921 | 729.5905 | 2.14 | SM d36:2 | [M+H]+ | C41H81N2O6P |
| 731.6072 | 731.6062 | 1.47 | SM d36:1 | [M+H]+ | C41H83N2O6P |
| 732.5550 | 732.5538 | 1.62 | PC 32:1 | [M+H]+ | C40H78NO8P |
| 734.5697 | 734.5694 | 0.30 | PC 32:0 | [M+H]+ | C40H80NO8P |
| 742.5373 | 742.5357 | 2.08 | PE 34:0 | [M+Na]+ | C39H78NO8PNa |
| 743.4631 | 743.4622 | 1.14 | PA 38:6 | [M+Na]+ | C41H69O8PNa |
| 745.4786 | 745.4779 | 0.96 | PA 38:5 | [M+Na]+ | C41H71O8PNa |
| 746.5704 | 746.5694 | 1.34 | PE 36:1 | [M+H]+ | C41H80NO8P |
| 746.6069 | 746.6058 | 1.48 | PC O-34:1 | [M+H]+ | C42H84NO7P |
| 748.5864 | 748.5851 | 1.77 | PE 36:0 | [M+H]+ | C41H82NO8P |
| 749.5104 | 749.5092 | 1.58 | PA 38:3 | [M+Na]+ | C41H75O8PNa |
| 751.5257 | 751.5248 | 1.22 | PA 38:2 | [M+Na]+ | C41H77O8PNa |
| 751.5708 | 751.5724 | -2.14 | SM d36:2 | [M+Na]+ | C41H81N2O6PNa |
| 753.5889 | 753.5881 | 1.08 | SM d36:1 | [M+Na]+ | C41H83N2O6PNa |
| 754.5370 | 754.5357 | 1.64 | PC 32:1 | [M+Na]+ | C40H78NO8PNa |
| 755.4999 | 755.4986 | 1.69 | PA O-40:7 | [M+Na]+ | C43H73O7PNa |

| | | | | | |
|---|---|---|---|---|---|
| 756.5499 | 756.5514 | -1.99 | PC 32:0 | [M+Na]+ | C40H80NO8PNa |
| 758.5704 | 758.5694 | 1.26 | PC 34:2 | [M+H]+ | C42H80NO8P |
| 759.6366 | 759.6375 | -1.16 | SM d38:1 | [M+H]+ | C43H87N2O6P |
| 760.5854 | 760.5851 | 0.41 | PC 34:1 | [M+H]+ | C42H82NO8P |
| 762.6013 | 762.6007 | 0.71 | PC 34:0 | [M+H]+ | C42H84NO8P |
| 764.5192 | 764.5225 | -4.31 | PE 38:6 | [M+H]+ | C43H74NO8P |
| 766.5367 | 766.5381 | -1.83 | PE 38:5 | [M+H]+ | C43H76NO8P |
| 768.5521 | 768.5514 | 0.98 | PE 36:1 | [M+Na]+ | C41H80NO8PNa |
| 768.5885 | 768.5878 | 0.96 | PC O-34:1 | [M+Na]+ | C42H84NO7PNa |
| 769.4786 | 769.4779 | 0.91 | PA 40:7 | [M+Na]+ | C43H71O8PNa |
| 770.5106 | 770.5097 | 1.24 | PC 32:1 | [M+K]+ | C40H78NO8PK |
| 770.5684 | 770.5670 | 1.86 | PE 36:0 | [M+Na]+ | C41H82NO8PNa |
| 771.4948 | 771.4935 | 1.61 | PA 40:6 | [M+Na]+ | C43H73O8PNa |
| 772.5265 | 772.5253 | 1.51 | PC 32:0 | [M+K]+ | C40H80NO8PK |
| 773.5103 | 773.5092 | 1.45 | PA 40:5 | [M+Na]+ | C43H75O8PNa |
| 773.5297 | 773.5303 | -0.74 | PG 34:0 | [M+Na]+ | C40H79NaO10P |
| 774.6014 | 774.6007 | 0.88 | PE 38:1 | [M+H]+ | C43H84NO8P |
| 780.5495 | 780.5514 | -2.44 | PC 34:2 | [M+Na]+ | C42H80NO8PNa |
| 780.5524 | 780.5538 | -1.74 | PC 36:5 | [M+H]+ | C44H78NO8P |
| 781.6207 | 781.6194 | 1.62 | SM d38:1 | [M+Na]+ | C43H87N2O6PNa |
| 782.5679 | 782.5670 | 1.09 | PC 34:1 | [M+Na]+ | C42H82NO8PNa |
| 782.5679 | 782.5694 | -1.98 | PC 36:4 | [M+H]+ | C44H80NO8P |
| 784.5825 | 784.5827 | -0.19 | PC 34:0 | [M+Na]+ | C42H84NO8PNa |
| 784.5825 | 784.5851 | -3.25 | PC 36:3 | [M+H]+ | C44H83NO8P |
| 786.5049 | 786.5044 | 0.65 | PE 38:6 | [M+Na]+ | C43H74NO8PNa |
| 786.6007 | 786.6007 | -0.07 | PC 36:2 | [M+H]+ | C44H84NO8P |
| 788.6166 | 788.6164 | 0.29 | PC 36:1 | [M+H]+ | C44H86NO8P |
| 790.5140 | 790.5147 | -0.90 | PE O-38:5 | [M+K]+ | C43H78KNO7P |
| 790.5349 | 790.5357 | -1.03 | PE 38:4 | [M+Na]+ | C43H78NO8PNa |
| 790.5370 | 790.5381 | -1.46 | PE 40:7 | [M+H]+ | C45H76NO8P |
| 792.4926 | 792.4940 | -1.78 | PC 34:4 | [M+K]+ | C42H76NO8PK |
| 792.5298 | 792.5304 | -0.79 | PE O-38:4 | [M+K]+ | C43H80NO7PK |
| 793.4764 | 793.4779 | -1.94 | PA 42:9 | [M+Na]+ | C45H71O8PNa |
| 794.5078 | 794.5097 | -2.38 | PC 34:3 | [M+K]+ | C42H78NO8PK |
| 796.5227 | 796.5253 | -3.29 | PC 34:2 | [M+K]+ | C42H80NO8PK |
| 796.5828 | 796.5827 | 0.11 | PE 38:1 | [M+Na]+ | C43H84NO8PNa |
| 796.5828 | 796.5851 | -2.91 | PE 40:4 | [M+H]+ | C45H82NO8P |
| 797.5092 | 797.5093 | -0.11 | PG O-36:3 | [M+K]+ | C42H79O9PK |
| 798.5411 | 798.5410 | 0.15 | PC 34:1 | [M+K]+ | C42H82NO8PK |
| 800.5574 | 800.5566 | 0.93 | PC 34:0 | [M+K]+ | C42H84NO8PK |
| 802.6329 | 802.6320 | 1.09 | PE 40:1 | [M+H]+ | C45H88NO8P |
| 804.5516 | 804.5514 | 0.28 | PC 36:4 | [M+Na]+ | C44H80NO8PNa |
| 806.5686 | 806.5694 | -1.02 | PC 38:6 | [M+H]+ | C46H80NO8P |

| | | | | | |
|---|---|---|---|---|---|
| 808.5823 | 808.5827 | -0.51 | PC 36:2 | [M+Na]+ | C44H84NO8PNa |
| 808.5849 | 808.5851 | -0.21 | PC 38:5 | [M+H]+ | C46H82NO8P |
| 810.5950 | 810.5983 | -4.08 | PC 36:1 | [M+Na]+ | C44H86NO8PNa |
| 810.5991 | 810.6007 | -2.00 | PC 38:4 | [M+H]+ | C46H84NO8P |
| 812.4957 | 812.4991 | -4.21 | PE O-40:8 | [M+K]+ | C45H76NO7PK |
| 812.5186 | 812.5202 | -1.95 | PS O-36:2 | [M+K]+ | C42H80NO9PK |
| 812.6174 | 812.6164 | 1.30 | PC 38:3 | [M+H]+ | C46H86NO8P |
| 813.6846 | 813.6844 | 0.26 | SM d42:2 | [M+H]+ | C47H93N2O6P |
| 814.5365 | 814.5359 | 0.81 | PC 34:1 OH | [M+K]+ | C42H82NO9PK |
| 816.5294 | 816.5304 | -1.26 | PE O-40:6 | [M+K]+ | C45H80NO7PK |
| 818.5077 | 818.5097 | -2.37 | PC 36:5 | [M+K]+ | C44H78NO8PK |
| 818.5448 | 818.5460 | -1.53 | PE O-40:5 | [M+K]+ | C45H82NO7PK |
| 820.5225 | 820.5253 | -3.39 | PC 36:4 | [M+K]+ | C44H80NO8PK |
| 820.5821 | 820.5851 | -3.61 | PE 42:6 | [M+H]+ | C47H82NO8P |
| 820.5851 | 820.5851 | -0.01 | PE 42:6 | [M+H]+ | C47H82NO8P |
| 824.6163 | 824.6164 | -0.06 | PE 42:4 | [M+H]+ | C47H86NO8P |
| 826.5732 | 826.5723 | 1.13 | PC 36:1 | [M+K]+ | C44H86NO8PK |
| 828.5517 | 828.5514 | 0.37 | PC 38:6 | [M+Na]+ | C46H80NO8PNa |
| 830.5543 | 830.5542 | 0.17 | PS 38:3 OH | [M+H]+ | C44H80NO11P |
| 830.5677 | 830.5670 | 0.79 | PC 38:5 | [M+Na]+ | C46H82NO8PNa |
| 830.5677 | 830.5694 | -2.11 | PC 40:8 | [M+H]+ | C48H80NO8P |
| 832.5790 | 832.5827 | -4.43 | PC 38:4 | [M+Na]+ | C46H84NO8PNa |
| 832.5831 | 832.5851 | -2.36 | PC 40:7 | [M+H]+ | C48H82NO8P |
| 834.6000 | 834.6007 | -0.94 | PC 40:6 | [M+H]+ | C48H84NO8P |
| 836.5168 | 836.5202 | -4.12 | PC 36:4 OH | [M+K]+ | C44H80NO9PK |
| 836.6161 | 836.6140 | 2.51 | PC 38:2 | [M+Na]+ | C46H88NO8PNa |
| 836.6161 | 836.6164 | -0.38 | PC 40:5 | [M+H]+ | C48H86NO8P |
| 836.6161 | 836.6140 | 2.49 | PC 38:2 | [M+Na]+ | C46H88NO8PNa |
| 838.6286 | 838.6296 | -1.27 | PC 38:1 | [M+Na]+ | C46H90NO8PNa |
| 838.6316 | 838.6320 | -0.54 | PC 40:4 | [M+H]+ | C48H88NO8P |
| 844.5259 | 844.5253 | 0.73 | PC 38:6 | [M+K]+ | C46H80NO8PK |
| 844.6799 | 844.6790 | 1.12 | PC 40:1 | [M+H]+ | C48H94NO8P |
| 848.5183 | 848.5201 | -2.17 | PC 40:10 | [M+Na]+ | C48H76NO8PNa |
| 850.6752 | 850.6743 | 1.10 | HexCer t42:1 | [M+Na]+ | C48H93NO9Na |
| 854.5663 | 854.5670 | -0.91 | PC 40:7 | [M+Na]+ | C48H82NO8PNa |
| 856.5814 | 856.5827 | -1.55 | PC 40:6 | [M+Na]+ | C48H84NO8PNa |
| 858.5245 | 858.5256 | -1.25 | PS 40:6 | [M+Na]+ | C46H78NO10PNa |
| 858.5852 | 858.5855 | -0.31 | PS 40:3 OH | [M+H]+ | C46H84NO11P |
| 872.7103 | 872.7103 | 0.01 | PC 42:1 | [M+H]+ | C50H98NO8P |

| Observed m/z | Theoretical m/z | mass error | species | ion | formula |
|---|---|---|---|---|---|
| 647.4656 | 647.4657 | -0.25 | PA 32:0 | [M-H]- | C35H69O8P |
| 671.4660 | 671.4657 | 0.43 | PA 34:2 | [M-H]- | C37H69O8P |
| 673.4813 | 673.4814 | -0.06 | PA 34:1 | [M-H]- | C37H71O8P |
| 675.4975 | 675.4970 | 0.71 | PA 34:0 | [M-H]- | C37H73O8P |
| 695.4660 | 695.4657 | 0.37 | PA 36:4 | [M-H]- | C39H69O8P |
| 697.4823 | 697.4814 | 1.31 | PA 36:3 | [M-H]- | C39H71O8P |
| 699.4971 | 699.4970 | 0.06 | PA 36:2 | [M-H]- | C39H73O8P |
| 700.5289 | 700.5287 | 0.27 | PE O-34:2 | [M-H]- | C39H76NO7P |
| 701.5132 | 701.5127 | 0.76 | PA 36:1 | [M-H]- | C39H75O8P |
| 714.5091 | 714.5079 | 1.70 | PE 34:2 | [M-H]- | C39H74NO8P |
| 716.5240 | 716.5236 | 0.55 | PE 34:1 | [M-H]- | C39H76NO8P |
| 718.5398 | 718.5392 | 0.82 | PE 34:0 | [M-H]- | C39H78NO8P |
| 719.4668 | 719.4657 | 1.54 | PA 38:6 | [M-H]- | C41H69O8P |
| 721.5031 | 721.5025 | 0.79 | PG 32:0 | [M-H]- | C38H75O10P |
| 723.4978 | 723.4970 | 1.12 | PA 38:4 | [M-H]- | C41H73O8P |
| 724.5299 | 724.5287 | 1.69 | PE O-36:4 | [M-H]- | C41H76NO7P |
| 725.5137 | 725.5127 | 1.37 | PA 38:3 | [M-H]- | C41H75O8P |
| 726.5445 | 726.5443 | 0.21 | PE O-36:3 | [M-H]- | C41H78NO7P |
| 727.5285 | 727.5283 | 0.29 | PA 38:2 | [M-H]- | C41H77O8P |
| 728.5606 | 728.5600 | 0.85 | PE O-36:2 | [M-H]- | C41H80NO7P |
| 729.5453 | 729.5440 | 1.75 | PA 38:1 | [M-H]- | C41H79O8P |
| 738.5086 | 738.5079 | 0.96 | PE 36:4 | [M-H]- | C41H74NO8P |
| 740.5250 | 740.5236 | 1.92 | PE 36:3 | [M-H]- | C41H76NO8P |
| 742.5393 | 742.5392 | 0.11 | PE 36:2 | [M-H]- | C41H78NO8P |
| 744.5551 | 744.5549 | 0.27 | PE 36:1 | [M-H]- | C41H80NO8P |
| 745.4827 | 745.4814 | 1.78 | PA 40:7 | [M-H]- | C43H71O8P |
| 745.5026 | 745.5025 | 0.10 | PG 34:2 | [M-H]- | C40H75O10P |
| 746.5143 | 746.5130 | 1.66 | PE P-38:6 | [M-H]- | C43H74NO7P |
| 746.5718 | 746.5705 | 1.74 | PE 36:0 | [M-H]- | C41H82NO8P |
| 747.4976 | 747.4970 | 0.75 | PA 40:6 | [M-H]- | C43H73O8P |
| 747.5191 | 747.5182 | 1.21 | PG 34:1 | [M-H]- | C40H77O10P |
| 748.5288 | 748.5287 | 0.18 | PE O-38:6 | [M-H]- | C43H76NO7P |
| 749.5129 | 749.5127 | 0.23 | PA 40:5 | [M-H]- | C43H75O8P |
| 750.5450 | 750.5443 | 0.94 | PE O-38:5 | [M-H]- | C43H78NO7P |
| 751.5297 | 751.5283 | 1.81 | PA 40:4 | [M-H]- | C43H77O8P |
| 752.5613 | 752.5600 | 1.83 | PE O-38:4 | [M-H]- | C43H80NO7P |
| 754.5397 | 754.5392 | 0.61 | PE 37:3 | [M-H]- | C42H78NO8P |
| 754.5759 | 754.5756 | 0.33 | PE O-38:3 | [M-H]- | C43H82NO7P |
| 755.5600 | 755.5596 | 0.49 | PA 40:2 | [M-H]- | C43H81O8P |
| 756.5920 | 756.5913 | 0.93 | PE O-38:2 | [M-H]- | C43H84NO7P |
| 757.5767 | 757.5753 | 1.90 | PA 40:1 | [M-H]- | C43H83O8P |
| 760.5140 | 760.5134 | 0.83 | PS 34:1 | [M-H]- | C40H76NO10P |

| | | | | | |
|---|---|---|---|---|---|
| 762.5081 | 762.5079 | 0.16 | PE 38:6 | [M-H]- | C43H74NO8P |
| 764.5248 | 764.5236 | 1.53 | PE 38:5 | [M-H]- | C43H76NO8P |
| 766.5397 | 766.5392 | 0.60 | PE 38:4 | [M-H]- | C43H78NO8P |
| 768.5551 | 768.5549 | 0.33 | PE 38:3 | [M-H]- | C43H80NO8P |
| 770.5714 | 770.5705 | 1.13 | PE 38:2 | [M-H]- | C43H82NO8P |
| 771.5185 | 771.5182 | 0.44 | PG 36:3 | [M-H]- | C42H77O10P |
| 772.5302 | 772.5287 | 1.98 | PE P-40:7 | [M-H]- | C45H76NO7P |
| 772.5871 | 772.5862 | 1.18 | PE 38:1 | [M-H]- | C43H84NO8P |
| 773.5139 | 773.5127 | 1.62 | PA 42:7 | [M-H]- | C45H75O8P |
| 773.5353 | 773.5338 | 1.98 | PG 36:2 | [M-H]- | C42H79O10P |
| 774.5445 | 774.5443 | 0.18 | PE P-40:6 | [M-H]- | C45H78NO7P |
| 776.4984 | 776.4988 | -0.51 | SHexCer d34:2 | [M-H]- | C40H75NO11S |
| 776.5603 | 776.5600 | 0.37 | PE O-40:6 | [M-H]- | C45H80NO7P |
| 778.5159 | 778.5145 | 1.84 | SHexCer d34:1 | [M-H]- | C40H77NO11S |
| 778.5407 | 778.5392 | 1.83 | PE 39:5 | [M-H]- | C44H78NO8P |
| 778.5769 | 778.5756 | 1.68 | PE O-40:5 | [M-H]- | C45H82NO7P |
| 782.6070 | 782.6069 | 0.08 | PE O-40:3 | [M-H]- | C45H86NO7P |
| 783.5912 | 783.5909 | 0.40 | PA 42:2 | [M-H]- | C45H85O8P |
| 784.6224 | 784.6226 | -0.24 | PE O-40:2 | [M-H]- | C45H88NO7P |
| 785.6078 | 785.6066 | 1.51 | PA 42:1 | [M-H]- | C45H87O8P |
| 786.5304 | 786.5291 | 1.76 | PS 36:2 | [M-H]- | C42H78NO10P |
| 788.5249 | 788.5236 | 1.70 | PE 40:7 | [M-H]- | C45H76NO8P |
| 788.5455 | 788.5447 | 1.00 | PS 36:1 | [M-H]- | C42H80NO10P |
| 790.5391 | 790.5392 | -0.20 | PE 40:6 | [M-H]- | C45H78NO8P |
| 792.5552 | 792.5549 | 0.39 | PE 40:5 | [M-H]- | C45H80NO8P |
| 794.5343 | 794.5341 | 0.23 | PS O-38:5 | [M-H]- | C44H78NO9P |
| 794.5719 | 794.5705 | 1.68 | PE 40:4 | [M-H]- | C45H82NO8P |
| 797.5340 | 797.5338 | 0.28 | PG 38:4 | [M-H]- | C44H79O10P |
| 798.6025 | 798.6018 | 0.79 | PE 40:2 | [M-H]- | C45H86NO8P |
| 798.6467 | 798.6465 | 0.30 | HexCer t40:1 | [M-H]- | C46H89NO9 |
| 802.5401 | 802.5392 | 1.02 | PE 41:7 | [M-H]- | C46H78NO8P |
| 804.5315 | 804.5301 | 1.78 | SHexCer d36:2 | [M-H]- | C42H79NO11S |
| 804.5563 | 804.5549 | 1.79 | PE 41:6 | [M-H]- | C46H80NO8P |
| 804.5929 | 804.5913 | 2.07 | PE O-42:6 | [M-H]- | C47H84NO7P |
| 806.5340 | 806.5341 | -0.14 | PE 40:6 OH | [M-H]- | C45H78NO9P |
| 806.5463 | 806.5458 | 0.72 | SHexCer d36:1 | [M-H]- | C42H81NO11S |
| 808.5111 | 808.5134 | -2.85 | PS 38:5 | [M-H]- | C44H76NO10P |
| 808.5613 | 808.5614 | -0.13 | SHexCer d36:0 | [M-H]- | C42H83NO11S |

| | | | | | |
|---|---|---|---|---|---|
| 808.6670 | 808.6672 | -0.20 | HexCer d42:2 | [M-H]- | C48H91NO8 |
| 809.5182 | 809.5186 | -0.39 | PI 32:0 | [M-H]- | C41H79O13P |
| 810.5266 | 810.5291 | -3.09 | PS 38:4 | [M-H]- | C44H78NO10P |
| 812.6631 | 812.6621 | 1.20 | HexCer t41:1 | [M-H]- | C47H91NO9 |
| 814.5618 | 814.5604 | 1.77 | PS 38:2 | [M-H]- | C44H82NO10P |
| 816.5561 | 816.5549 | 1.47 | PE 42:7 | [M-H]- | C47H80NO8P |
| 816.5772 | 816.5760 | 1.40 | PS 38:1 | [M-H]- | C44H84NO10P |
| 818.5342 | 818.5341 | 0.06 | PS P-40:6 | [M-H]- | C46H78NO9P |
| 818.5705 | 818.5705 | -0.08 | PE 42:6 | [M-H]- | C47H82NO8P |
| 819.5182 | 819.5182 | -0.01 | PG 40:7 | [M-H]- | C46H77O10P |
| 820.5244 | 820.5250 | -0.77 | SHexCer t36:2 | [M-H]- | C42H79NO12S |
| 822.5410 | 822.5407 | 0.37 | SHexCer t36:1 | [M-H]- | C42H81NO12S |
| 824.6632 | 824.6621 | 1.31 | HexCer t42:2 | [M-H]- | C48H91NO9 |
| 826.6786 | 826.6778 | 1.04 | HexCer t42:1 | [M-H]- | C48H93NO9 |
| 832.5105 | 832.5134 | -3.44 | PS 40:7 | [M-H]- | C46H76NO10P |
| 832.5610 | 832.5614 | -0.46 | SHexCer d38:2 | [M-H]- | C44H83NO11S |
| 834.5068 | 834.5079 | -1.29 | PE 44:12 | [M-H]- | C49H74NO8P |
| 834.5291 | 834.5291 | 0.07 | PS 40:6 | [M-H]- | C46H78NO10P |
| 834.5775 | 834.5771 | 0.48 | SHexCer d38:1 | [M-H]- | C44H85NO11S |
| 835.5336 | 835.5342 | -0.67 | PI 34:1 | [M-H]- | C43H81O13P |
| 836.5460 | 836.5447 | 1.52 | PS 40:5 | [M-H]- | C46H80NO10P |
| 836.5938 | 836.5927 | 1.27 | SHexCer d38:0 | [M-H]- | C44H87NO11S |
| 837.5514 | 837.5499 | 1.79 | PI 34:0 | [M-H]- | C43H83O13P |
| 838.5405 | 838.5392 | 1.53 | PE 44:10 | [M-H]- | C49H78NO8P |
| 838.5616 | 838.5604 | 1.50 | PS 40:4 | [M-H]- | C46H82NO10P |
| 842.5915 | 842.5917 | -0.18 | PS 40:2 | [M-H]- | C46H86NO10P |
| 844.6064 | 844.6073 | -1.05 | PS 40:1 | [M-H]- | C46H88NO10P |
| 848.5568 | 848.5563 | 0.55 | SHexCer t38:2 | [M-H]- | C44H83NO12S |
| 850.5729 | 850.5720 | 1.07 | SHexCer t38:1 | [M-H]- | C44H85NO12S |
| 856.5105 | 856.5134 | -3.36 | PS 42:9 | [M-H]- | C48H76NO10P |
| 857.5176 | 857.5186 | -1.11 | PI 36:4 | [M-H]- | C45H79O13P |
| 858.5838 | 858.5866 | -3.25 | PS 40:2 OH | [M-H]- | C46H86NO11P |
| 859.5333 | 859.5342 | -1.10 | PI 36:3 | [M-H]- | C45H81O13P |
| 860.5416 | 860.5447 | -3.67 | PS 42:7 | [M-H]- | C48H80NO10P |

| | | | | | |
|---|---|---|---|---|---|
| 860.5918 | 860.5927 | -1.05 | SHexCer d40:2 | [M-H]- | C46H87NO11S |
| 861.5492 | 861.5499 | -0.78 | PI 36:2 | [M-H]- | C45H83O13P |
| 862.5602 | 862.5604 | -0.22 | PS 42:6 | [M-H]- | C48H82NO10P |
| 862.6077 | 862.6084 | -0.81 | SHexCer d40:1 | [M-H]- | C46H89NO11S |
| 863.5653 | 863.5655 | -0.25 | PI 36:1 | [M-H]- | C45H85O13P |
| 864.5878 | 864.5876 | 0.24 | SHexCer t39:1 | [M-H]- | C45H87NO12S |
| 864.6242 | 864.6240 | 0.26 | SHexCer d40:0 | [M-H]- | C46H91NO11S |
| 865.5027 | 865.5025 | 0.17 | PG 44:12 | [M-H]- | C50H75O10P |
| 866.5894 | 866.5917 | -2.65 | PS 42:4 | [M-H]- | C48H86NO10P |
| 867.5396 | 867.5393 | 0.39 | PI O-38:6 | [M-H]- | C47H81O12P |
| 870.6233 | 870.6230 | 0.36 | PS 42:2 | [M-H]- | C48H90NO10P |
| 872.6384 | 872.6386 | -0.23 | PS 42:1 | [M-H]- | C48H92NO10P |
| 874.6078 | 874.6084 | -0.69 | SHexCer d41:2 | [M-H]- | C47H89NO11S |
| 876.5867 | 876.5876 | -1.04 | SHexCer t40:2 | [M-H]- | C46H87NO12S |
| 876.6231 | 876.6240 | -1.03 | SHexCer d41:1 | [M-H]- | C47H91NO11S |
| 878.6027 | 878.6033 | -0.67 | SHexCer t40:1 | [M-H]- | C46H89NO12S |
| 880.6188 | 880.6189 | -0.18 | SHexCer t40:0 | [M-H]- | C46H91NO12S |
| 881.5186 | 881.5186 | 0.08 | PI 38:6 | [M-H]- | C47H79O13P |
| 883.5356 | 883.5342 | 1.55 | PI 38:5 | [M-H]- | C47H81O13P |
| 885.5500 | 885.5499 | 0.14 | PI 38:4 | [M-H]- | C47H83O13P |
| 886.6095 | 886.6084 | 1.27 | SHexCer d42:3 | [M-H]- | C48H89NO11S |
| 887.5672 | 887.5655 | 1.86 | PI 38:3 | [M-H]- | C47H85O13P |
| 888.6242 | 888.6240 | 0.22 | SHexCer d42:2 | [M-H]- | C48H91NO11S |
| 890.6396 | 890.6397 | -0.12 | SHexCer d42:1 | [M-H]- | C48H93NO11S |
| 892.6194 | 892.6189 | 0.48 | SHexCer t41:1 | [M-H]- | C47H91NO12S |
| 892.6555 | 892.6553 | 0.26 | SHexCer d42:0 | [M-H]- | C48H95NO11S |
| 901.5457 | 901.5448 | 0.99 | PI 38:4 OH | [M-H]- | C47H83O14P |
| 902.6404 | 902.6397 | 0.87 | SHexCer d43:2 | [M-H]- | C49H93NO11S |
| 904.6190 | 904.6189 | 0.11 | SHexCer t42:2 | [M-H]- | C48H91NO12S |

| | | | | | |
|---|---|---|---|---|---|
| 904.6567 | 904.6553 | 1.53 | SHexCer d43:1 | [M-H]- | C49H95NO11S |
| 906.6348 | 906.6346 | 0.23 | SHexCer t42:1 | [M-H]- | C48H93NO12S |
| 907.5344 | 907.5342 | 0.16 | PI 40:7 | [M-H]- | C49H81O13P |
| 908.6503 | 908.6502 | 0.11 | SHexCer t42:0 | [M-H]- | C48H95NO12S |
| 909.5509 | 909.5499 | 1.16 | PI 40:6 | [M-H]- | C49H83O13P |
| 909.5509 | 909.5499 | 1.11 | PI 40:6 | [M-H]- | C49H83O13P |
| 911.5659 | 911.5655 | 0.45 | PI 40:5 | [M-H]- | C49H85O13P |
| 913.5813 | 913.5812 | 0.19 | PI 40:4 | [M-H]- | C49H87O13P |
| 914.6395 | 914.6397 | -0.13 | SHexCer d44:3 | [M-H]- | C50H93NO11S |
| 916.6583 | 916.6553 | 3.28 | SHexCer d44:2 | [M-H]- | C50H95NO11S |
| 918.6737 | 918.6710 | 3.03 | SHexCer d44:1 | [M-H]- | C50H97NO11S |
| 920.6520 | 920.6502 | 1.88 | SHexCer t43:1 | [M-H]- | C49H95NO12S |
| 932.6472 | 932.6502 | -3.21 | SHexCer t44:2 | [M-H]- | C50H95NO12S |
| 934.6631 | 934.6659 | -2.96 | SHexCer t44:1 | [M-H]- | C50H97NO12S |
| 953.5180 | 953.5186 | -0.63 | PI 44:12 | [M-H]- | C53H79O13P |

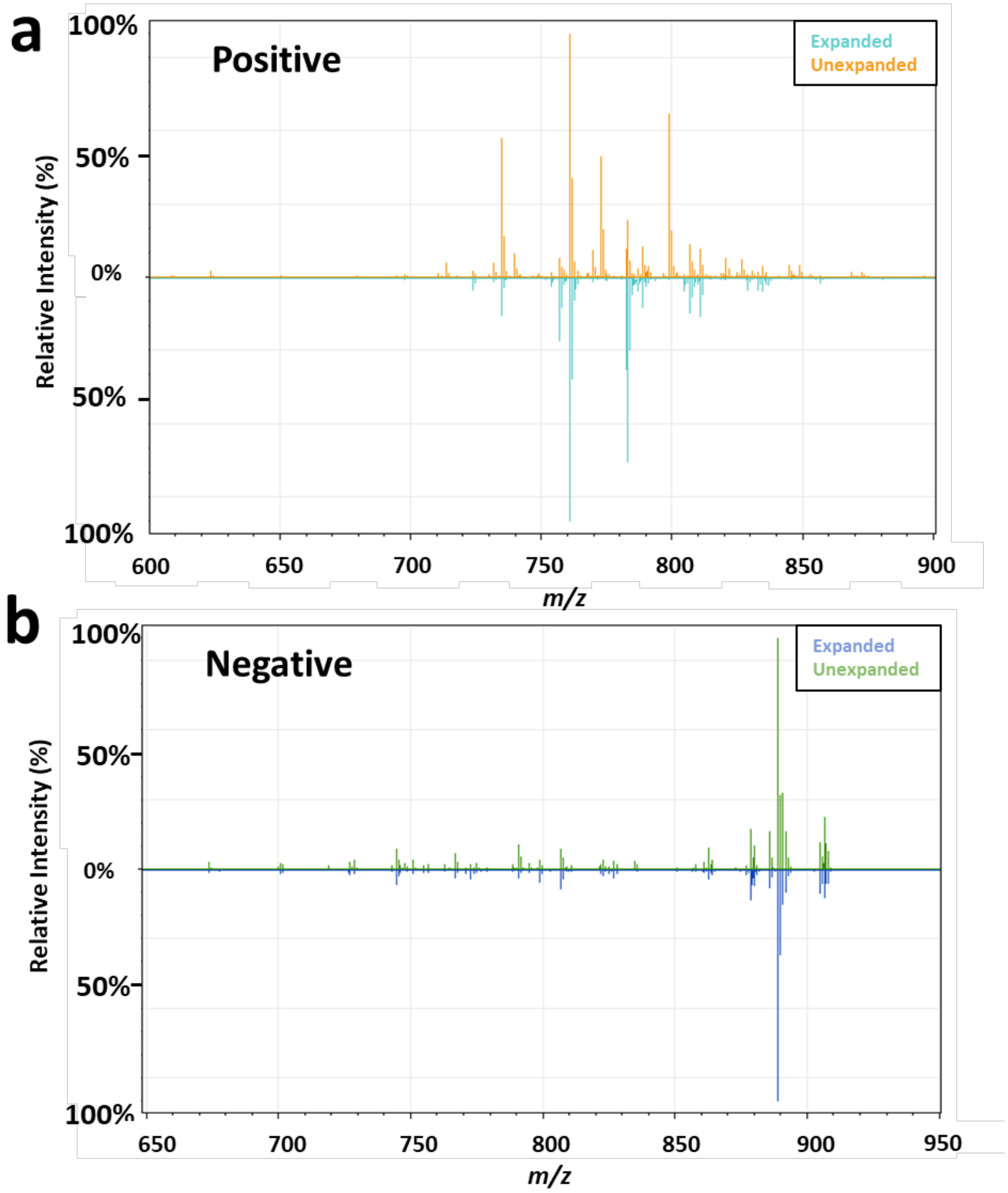

**Supplemental Figure S1:** Butterfly spectra of expanded versus unexpanded tissue in (a) positive and (b) negative ion modes.

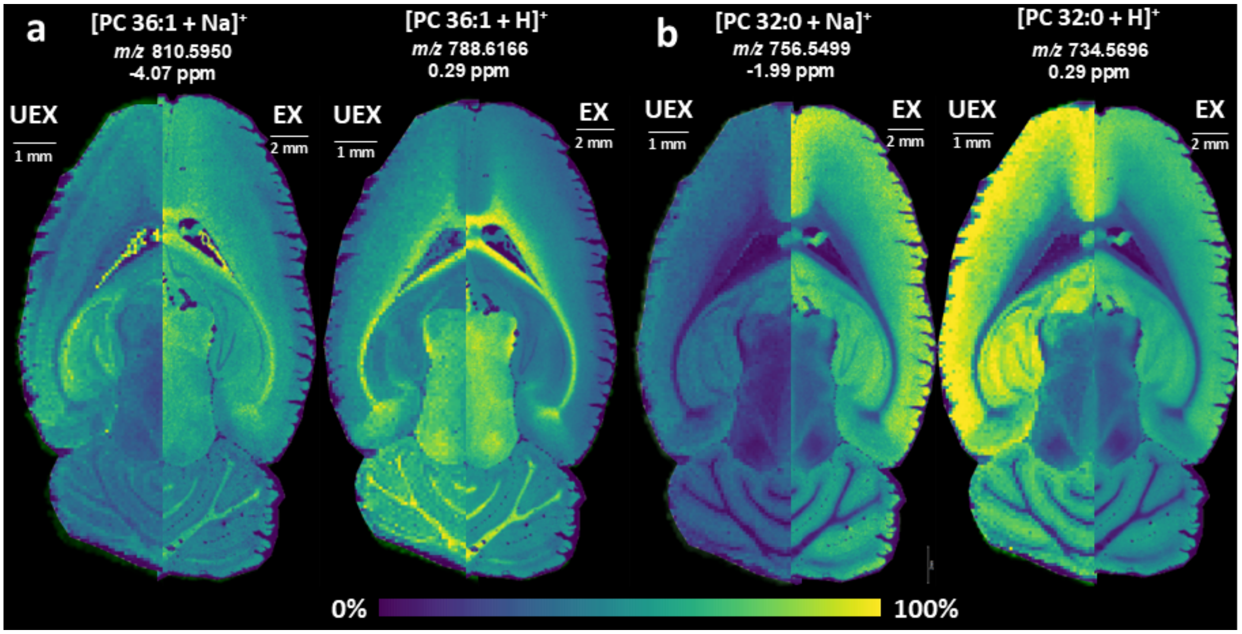

**Supplemental Figure S2:** Spatial distributions of sodiated and protonated lipid ion types are compared between unexpanded (left) and expanded (right) tissues. (a) Similar distributions are observed for PC 36:1 for the sodiated ion type in the expanded tissue and the protonated ion type in both the expanded and unexpanded tissues, which all differ from the sodiated ion type in the unexpanded sample. (b) Conversely, similar distributions are observed for protonated and sodiated ion types of PC 32:0 in both unexpanded and expanded tissue. Ion images for expanded and unexpanded tissues are set to the same intensity scale.

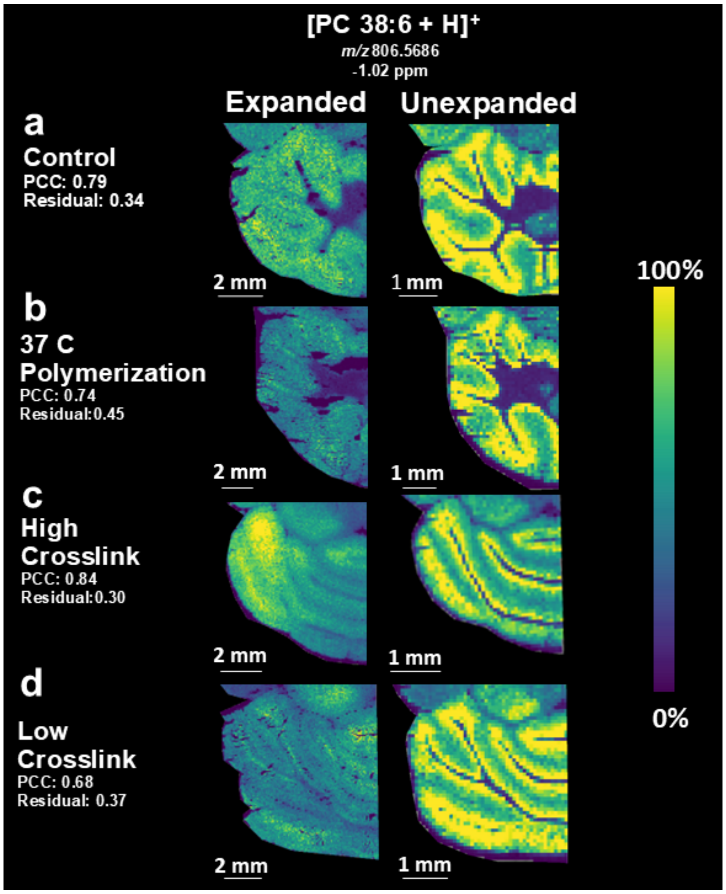

**Supplemental Figure S3*:*** ExIMS of mouse brain cerebellum is performed using variations in the expansion procedure, including (a) polymerization at room temperature, (b) polymerization at 37 °C, (c) expansion with a higher crosslinker density hydrogel, and (d) expansion with a lower crosslinker density hydrogel. Imaging mass spectrometry ion images of [PC 38:6 + H]$^+$ are displayed for expanded (left) and unexpanded (right) serial tissue sections. The expanded images all show higher intensity ion signal in the granular. The higher density crosslinker results in the most similar spatial distribution compared to the unexpanded tissue.

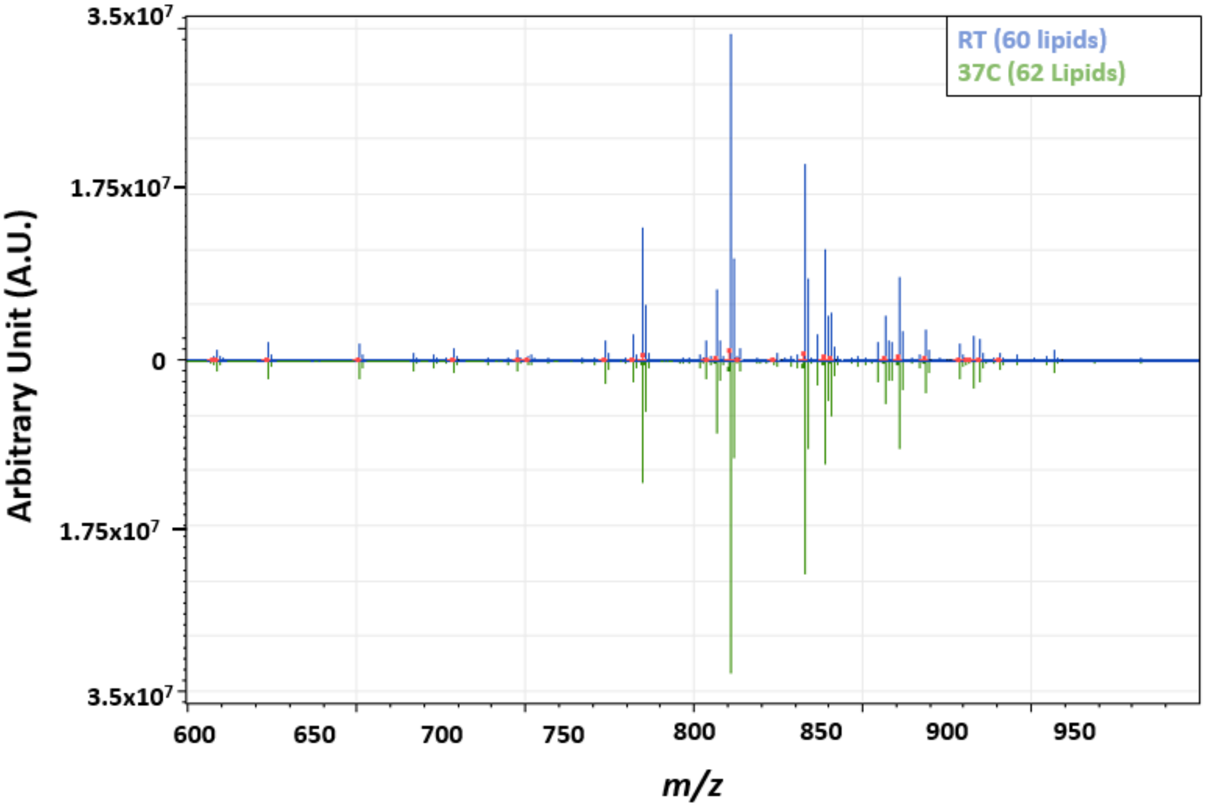

**Supplemental Figure S4:** Butterfly positive ion mode spectra from expanded tissue polymerized at room temperature for four hours versus 37C for two hours.
